# Harmonisation of multi-site MRS data with ComBat

**DOI:** 10.1101/2021.08.04.455132

**Authors:** Tiffany K. Bell, Kate J. Godfrey, Ashley L. Ware, Keith Owen Yeates, Ashley D. Harris

## Abstract

Magnetic resonance spectroscopy (MRS) is a non-invasive neuroimaging technique used to measure brain chemistry *in vivo* and has been used to study the healthy brain as well as neuropathology in numerous neurological disorders. The number of multi-site studies using MRS are increasing; however, non-biological variability introduced during data collection across multiple sites, such as differences in scanner vendors and site-specific acquisition implementations for MRS, can obscure detection of biological effects of interest. ComBat is a data harmonisation technique that can remove non-biological sources of variance in multisite studies. It has been validated for use with structural and functional MRI metrics but not for MRS metabolites. This study investigated the validity of using ComBat to harmonize MRS metabolites for vendor and site differences. Analyses were performed using data acquired across 20 sites and included edited MRS for GABA+ (N=218) and macromolecule-suppressed GABA data (N=209), as well as standard PRESS data to quantify NAA, creatine, choline, and glutamate (N=195). ComBat harmonisation successfully mitigated vendor and site differences for all metabolites of interest. Moreover, significant associations were detected between sex and choline levels and between age and glutamate and GABA+ levels that were not detectable prior to harmonisation, confirming the importance of removing site and vendor effects in multi-site data. In conclusion, ComBat harmonisation can be successfully applied to MRS data in multi-site MRS studies.

**Highlights:**

- Multi-site MRS data contains sources of non-biological variance that can mask biological effects of interest.
- ComBat data harmonisation successfully removes variance contributed by scanner and site differences.
- Removal of this variance revealed biological effects that were previously not detected.
- ComBat harmonisation can be successfully applied to MRS data in multi-site MRS studies

## INTRODUCTION

Magnetic resonance spectroscopy (MRS) enables non-invasive *in vivo* detection of brain metabolite levels. Thus, it is a useful technique to assess brain chemistry in healthy brain function, as well as changes in brain chemistry in neurological and neuropsychiatric disorders.

As with all neuroimaging modalities, multi-site MRS studies are becoming increasingly common to increase participant recruitment and statistical power to detect group differences, longitudinal changes, and relationships among brain chemistry and neuropsychological outcomes. Multi-site studies are particularly advantageous for assessing changes in brain chemistry, however, differences in MRS sequence implementations across vendors and sites can complicate data aggregation. For example, MRS sequence parameters such as pulse timings and pulse profiles differ between vendors, and these differences can lead to differences in signal (Near et al., 2013). Mikkelsen et al. (2017) recently conducted a study specifically aiming to examine site differences in MRS data. Twenty-four research sites collected GABA-edited and standard point-resolved spectroscopy (PRESS) MRS measures that resulted in data that was acquired using three different scanner vendors, seven different scanner models, five different shimming methods and four different water suppression techniques. Moreover, multiple variants of the MRS sequences themselves were used. Additionally, MRS data analysis is further complicated by the need for an internal reference signal, typically creatine or water (Near et al., 2020). The need for a separate water acquisition when referencing to water, as well as differences in the structural scan that is used for voxel segmentation are factors that may further increase non-biological (site or vendor) related variability. This variance is very typical of multi-site studies and will introduce a large amount of non-biological variability that may obscure the detection of biological effects of interest.

Indeed, Mikkelsen et al. showed significant effects of both vendor and site on GABA+ and a significant effect of site on macromolecule-suppressed (MM-sup) GABA levels both when referencing to creatine (Mikkelsen et al., 2017) and water (Mikkelsen et al., 2019). Považan et al., (2020) found similar effects of vendor and site on multiple metabolites quantified from PRESS data and referenced to creatine. However, none of these studies found significant effects of age on metabolite levels. This is perhaps surprising, as multiple studies have shown relationships between metabolite levels and age (see Cichocka and Bereś, 2018 for a systematic review on metabolite changes and (Porges et al., 2021) for a meta-analysis on GABA+ changes throughout the lifespan). The absence of age effects may therefore be due to confounding effects of vendor and/or site (Fortin et al., 2017).

Data harmonisation methods aim to remove unwanted variability caused by site or vendor differences while preserving true biological variability within the measures. In particular, ComBat, an empirical Bayesian method for data harmonisation originally developed for harmonisation of gene expression data (Johnson et al., 2007), has shown greater effectiveness at removing this unwanted variance in diffusion tensor imaging (DTI) data than other methods of harmonisation, including global scaling and functional normalisation (Fortin et al., 2017). In addition to DTI data, ComBat has been successfully applied to remove variance due to non-biological variables of interest in both cortical thickness (Fortin et al., 2018) and functional connectivity data (Yu et al., 2018). Further, the removal of non-biological variance using ComBat increased statistical power for detecting differences in brain structure between individuals with schizophrenia and healthy individuals compared to a random effects meta-analysis (Radua et al., 2020).

Despite the promising utility of ComBat for harmonisation of multisite structural and functional MRI data, it is yet to be validated with MRS data. This study aimed to validate the use of ComBat harmonisation on single voxel MRS data in healthy adults. Linear models were used to measure the variance contributed by vendor and site to metabolite levels both prior to and after harmonisation. Additionally, to examine whether ComBat harmonisation maintains true biological effects, age- and sex-related variance in metabolite levels was investigated both prior to and after harmonisation.

## METHODS

### Dataset

Data were obtained from the NITRIC Big GABA repository (www.nitrc.org/projects/biggaba/), which consists of single voxel MRS data acquired on 3T MRI scanners from the three major vendors, General Electric (GE), Philips, and Siemens (see Mikkelsen et al., 2017 for details on data acquisition). Scanning was conducted in accordance with ethical standards set by the institutional review board (IRB) at each site, including the sharing of anonymized data. Briefly, data were collected from participants aged 18-35 from a 3×3×3 cm^3^ voxel in the parietal lobe using a standard GABA+ edited sequence (TR/TE = 2000/68 ms, 320 averages, 15 ms editing pulses at 1.9 and 7.46 ppm), a MM-sup GABA edited sequence (TR/TE = 2000/80 ms for GE and Phillips, TR/TE = 2000/68ms for Siemens, 320 averages, 20 ms editing pulses at 1.9 and 1.5 ppm), and a standard PRESS sequence (TR/TE = 2000/35 ms, 64 averages). MEGA-PRESS data (both GABA+ and macromolecule suppressed, MM-sup GABA) were included from 20 sites and PRESS data were included from 19 sites (Table 1).

**Table 1:**
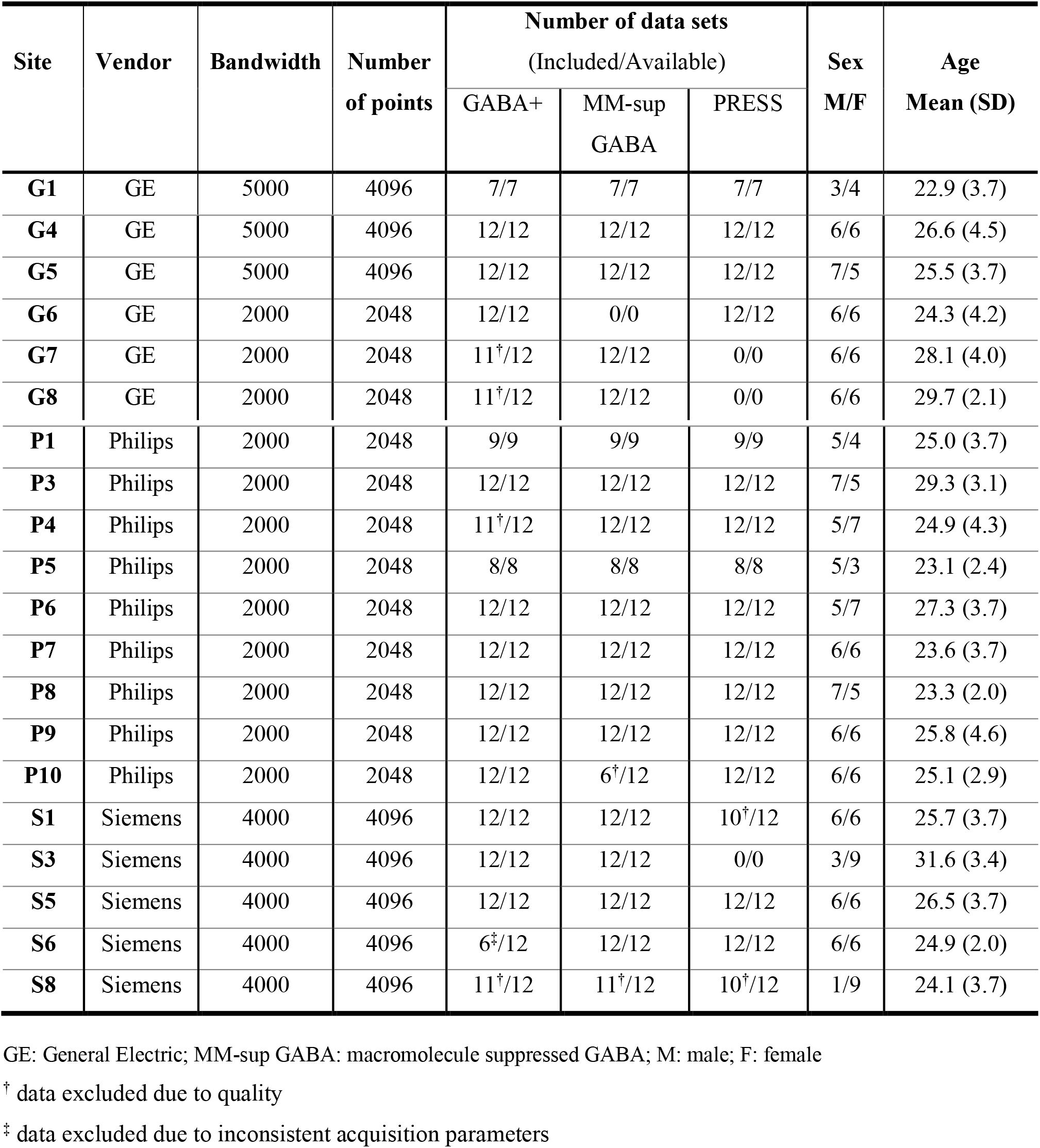
Summary of acquisition parameters that varied between sites, number of datasets available in the repository for each site and the number of datasets included in this analysis and participant demographics.

### Data Quantification

MEGA-PRESS data (both GABA+ and MM-sup GABA) were preprocessed and quantified with Gannet 3.2 (Edden et al., 2014) using the software’s automated analysis pipeline specified in the commands ‘GannetLoad’ and ‘GannetFit’. This procedure includes coil combination, removal of bad averages, and frequency and phase correction with spectral registration and modelling of the 3.0 ppm GABA+ or MM-sup GABA peak with a 5 parameter Gaussian function and modelling the unsuppressed water peak with a 7 parameter Voigt function. Coil combination for GE and Siemens PRESS data was performed in FID-A (Simpson et al., 2017), coil combination was performed on Philips scanners prior to export. Data were quantified using LCModel (Provencher, 2001). Generic basis sets were simulated rather than simulating basis sets based on individual acquisition pulse waveforms to increase the site-specific variance in quantified metabolite values (Považan et al., 2020). Basis sets were simulated using FID-A (te1=18 ms, te2=17 ms, number of points and spectral width were based on the acquired data) and included the following metabolites: alanine, aspartate, glycerophospocholine, phosphocholine, creatine, phosphocreatine, GABA, glutamate, glutamine, lactate, myo-inositol, n-acetylaspartate (NAA), n-acetylaspartylglutamate (NAAG), scyllo-inositol, glutathione, glucose and taurine. LCModel analysis used the default parameters (DOECC = T, DOWS = T, WCONC = 35880, ATTH20 = 0.7, ATTMET = 1.0). The LCModel quantified metabolites included in the harmonisation analysis were tNAA (total NAA, NAA+NAAG), tCr (total creatine, creatine + phosphocreatine), tCho (total choline, glycerophosphocholine + phosphocholine), Glx (glutamate + glutamine) and Myo-inositol (Myo-Ins).

Data were quantified as molal units and corrected for partial volume effects using the equation specified in Near et al. (2020). MEGA-PRESS data quantification included the α-correction, which assumes twice as much GABA in grey matter as in white matter (Harris et al., 2015). A secondary, confirmatory analysis for all metabolites (except tCr) was performed using tCr-referenced data.

### Harmonisation

Data were harmonised using the neuroComBat function version 1.0.5 (available at https://github.com/Jfortin1/ComBatHarmonization/tree/master/R) implemented in R version 4.0.3 (R Core Team (2020) https://www.R-project.org/). ComBat harmonisation corrects for one covariate while maintaining the variance from other covariates by estimating an empirical statistical distribution for each parameter. To do this, ComBat assumes all data points share the same common distribution, and therefore uses information from each data point to inform the statistical properties of the site and vendor effects (Fortin et al., 2017). Combat was used to correct for individual site (n=20 MEGA-PRESS/n=19 PRESS) and vendor (n=3) variance for each metabolite separately (as the distribution of one metabolite is not expected to be related to the distribution of another) and therefore used without empirical Bayes as the number of features was smaller than the number of participants. A matrix including the biological covariates of interest (grey matter fraction (fGM/(fGM+fWM)), age and sex) was included to preserve biological variability.

### Statistical Analysis

We followed the linear model procedure specified in Mikkelsen et al. (2017) to evaluate the presence of significant non-biological (vendor and site) and biological (grey matter fraction, age, and sex) effects in both the original and harmonised data. If vendor or site effects were significant in the data prior to harmonisation, but not following harmonisation, we considered the harmonisation procedure successful.

Vendor and site level effects on each metabolite were assessed separately using a three-level unconditional linear mixed-effects model, implemented using the R package lme4 version 1.1-26 (D et al., 2015), with vendor and site included as random effects. To validate that ComBat harmonisation maintains true biological effects, secondary multilevel analyses were performed to test for effects of grey-matter fraction, age, and sex on metabolite levels (included as fixed effects). Goodness of fit was calculated as a log-likelihood statistic. To assess the significance of each effect, likelihood ratio tests were performed by comparing a model with the effect of interest to a second model without the effect of interest. A p*-*value ≤ 0.05 was considered the threshold for significance. If an effect was significant, it was retained in the next assessed model for that metabolite; if not significant, it was removed. Effects were assessed in the following order: vendor, site, grey matter fraction, age, sex.

## RESULTS

### Evaluation of ComBat Harmonisation

#### Water referenced data

Prior to harmonization, vendor effects were significant for all metabolites and site effects were significant for all metabolites except MM-sup GABA. Given that both vendor and site were significantly associated with metabolite levels, and vendor variance was expected to be nested within site variance, data were harmonised by site. After harmonisation by site, neither vendor nor site effects were significantly associated with any metabolite. See Figure 1 for data visualisation prior to and after harmonisation, Table 2 for summaries of metabolite effects pre- and post-harmonisation, and supplementary material for model values.

**Figure 1:**
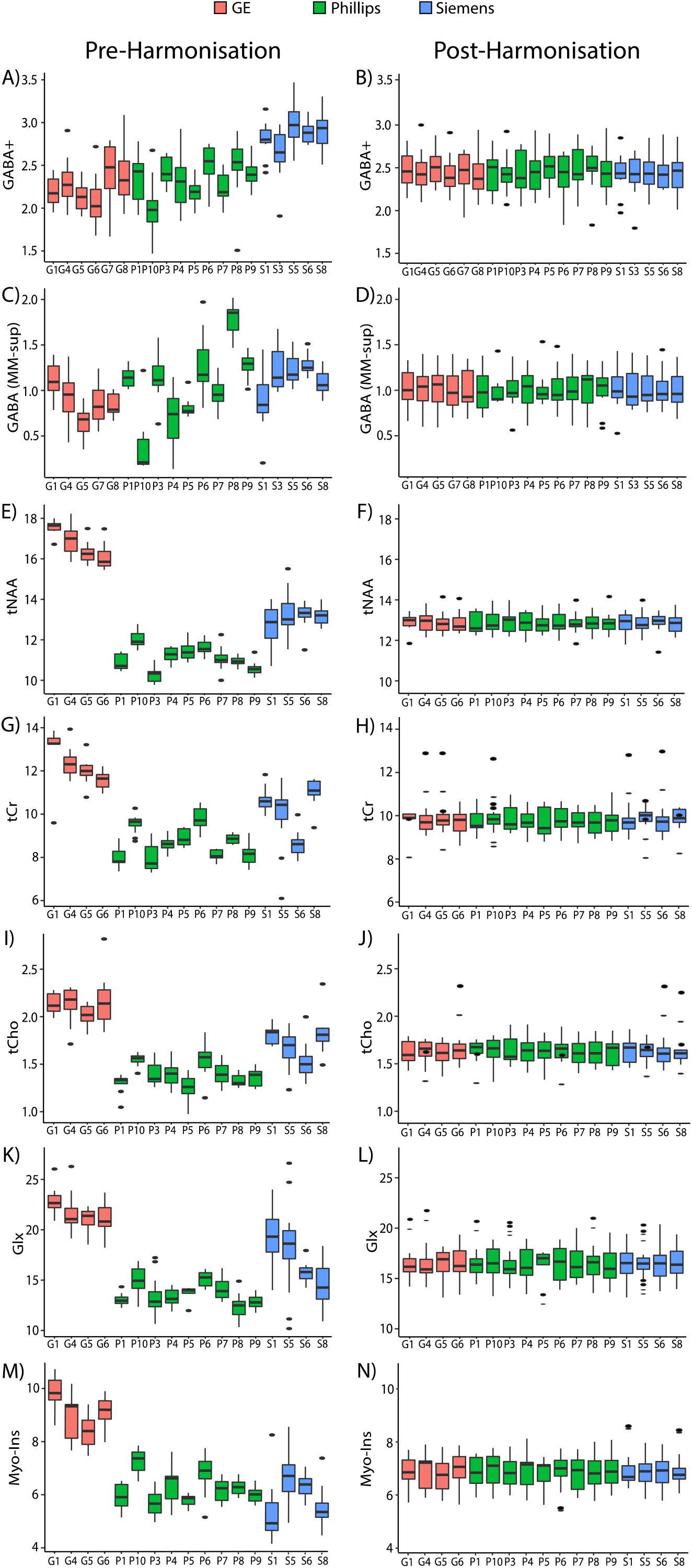
Metabolite levels pre and post-harmonisation. Box plots show median and interquartile range (IQR), dots represent outliers (more that 1.5xIQR). (A) GABA+ levels prior to harmonisation. (B) GABA+ levels following harmonisation. (C) MM-sup GABA levels prior to harmonisation. (D) MM-sup GABA levels following harmonisation. (E) tNAA levels prior to harmonisation. (F) tNAA levels following harmonisation. (G) tCr levels prior to harmonisation. (H) tCr levels following harmonisation. (I) tCho levels prior to harmonisation. (J) tCho levels following harmonisation. (K) Glx levels prior to harmonisation. (L) Glx levels following harmonisation. (M) Myo-Ins levels prior to harmonisation. (N) Myo-Ins levels following harmonisation. GABA+: GABA + macromolecules; MM-sup GABA: macromolecule suppressed GABA. (tNAA: total NAA (NAA+NAAG); tCr: total creatine (creatine + phosphocreatine); tCho: total choline (glycerophosphocholine + phosphocholine); Glx: glutamate + glutamine; Myo-Ins: Myo-inositol.

**Table 2:** Summary of significant model effects on water referenced metabolite data prior to and following harmonisation

|  |  | GABA+ | MM-sup GABA | tNAA | tCr | tCho | Glx | Myo |
| --- | --- | --- | --- | --- | --- | --- | --- | --- |
| **Pre-harmonisation** | Vendor | ✓ | ✓ | ✓ | ✓ | ✓ | ✓ | ✓ |
|  | Site | ✓ |  | ✓ | ✓ | ✓ | ✓ | ✓ |
|  | GMF |  |  |  |  |  |  |  |
|  | Age |  |  |  |  |  |  |  |
|  | Sex |  |  |  |  |  |  |  |
| **Post-harmonisation** | Vendor |  |  |  |  |  |  |  |
|  | Site |  |  |  |  |  |  |  |
|  | GMF | ✓ |  |  |  |  |  |  |
|  | Age |  |  |  |  |  |  |  |
|  | Sex |  |  |  |  | ✓ |  |  |

#### Creatine referenced data

Prior to harmonisation, vendor, but not site, was significantly associated with all creatine referenced metabolites. Thus, data were initially harmonised by vendor. However, the effect of vendor remained significant in the model, so data was therefore harmonised by site. After harmonisation by site, no vendor or site effects were significant (Table 3, see supplementary material for model values).

**Table 3:** Summary of significant model effects on creatine referenced metabolite data prior to and following harmonisation

|  |  | GABA+/tCr | MM-sup<br>GABA/tCr | tNAA/tCr | tCr/tCr | tCho/tCr | Glx/tCr | Myo/tCr |
| --- | --- | --- | --- | --- | --- | --- | --- | --- |
| **Pre-harmonisation** | Vendor | ✓ | ✓ | ✓ | ✓ | ✓ | ✓ | ✓ |
|  | Site |  |  |  |  |  |  |  |
|  | GMF |  |  |  |  |  |  |  |
|  | Age |  |  |  |  |  |  |  |
|  | Sex |  |  |  |  | ✓ |  |  |
| **Post-harmonisation** | Vendor |  |  |  |  |  |  |  |
|  | Site |  |  |  |  |  |  |  |
|  | GMF |  |  |  |  |  |  |  |
|  | Age | ✓ |  |  |  |  | ✓ |  |
|  | Sex |  |  |  |  | ✓ |  |  |

✓ denotes a significant contribution to the model
GMF: grey matter fraction; GABA+: GABA + macromolecules; MM-sup GABA: macromolecules suppressed GABA; tNAA: total n-acetylaspartate; tCr: total creatine; tCho: total choline; Glx: glutamate + glutamine; Myo: Myo-inositol.

### Biological Variability

#### Water referenced data

Prior to harmonisation, grey matter fraction, age, and sex were not significantly associated with water referenced metabolite levels. Following harmonisation, grey matter fraction was significantly associated with GABA+ levels and sex was significantly associated with tCho levels, with males having higher tCho levels than females (Figure 3, Table 2), see supplementary material for statistical model values.

**Figure 2.**
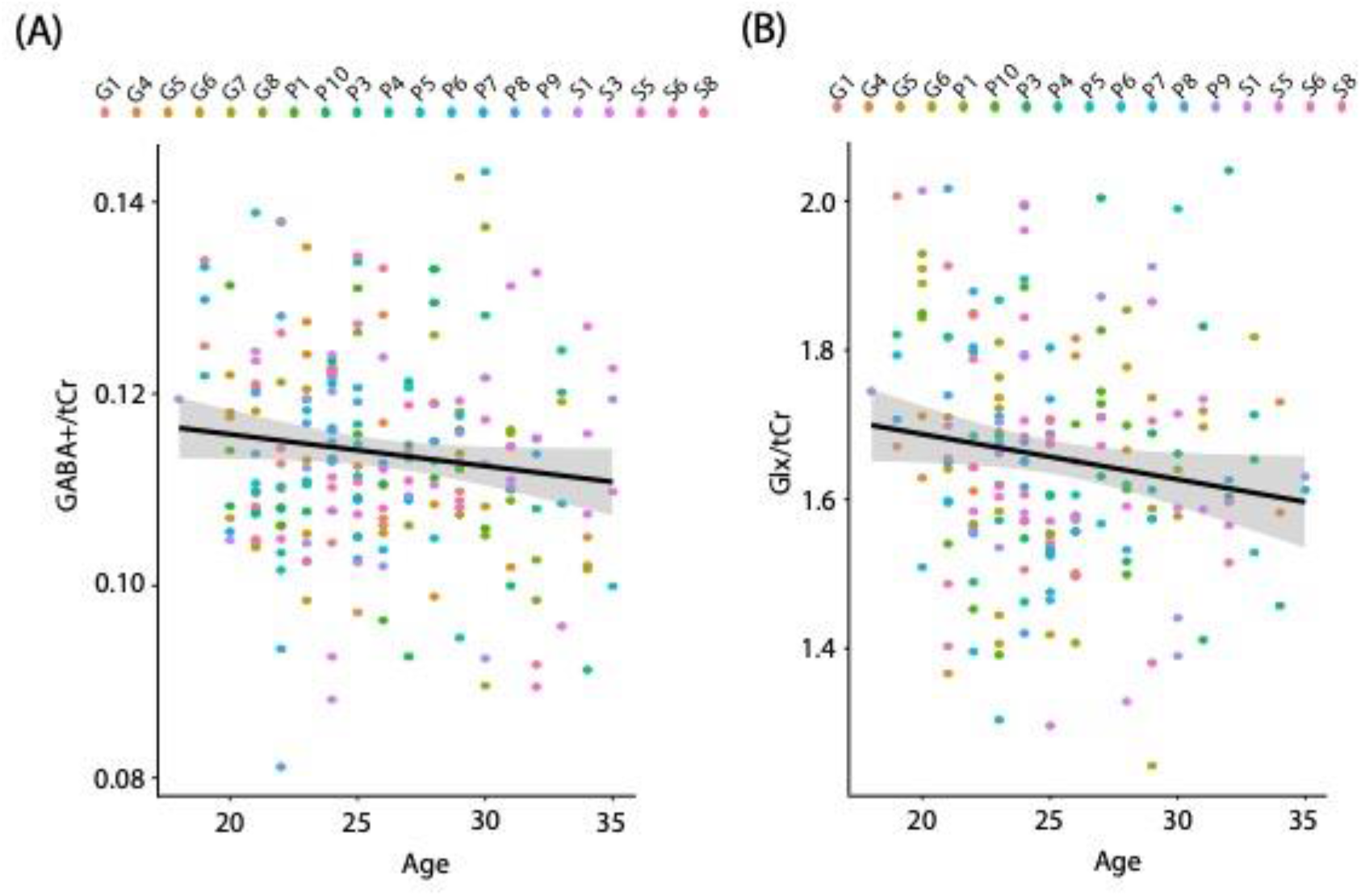
(A) Negative relationship between GABA+/tCr and age following harmonisation; (B) Negative relationship between Glx/tCr and age following harmonisation. Grey shading indiciates 95% confidence intervals, each dot represents a single individual, with dots colour coded by site. GABA+: GABA+ macromolecules; tCr: total creatine; Glx: glutamate + glutamine.

**Figure 3:**
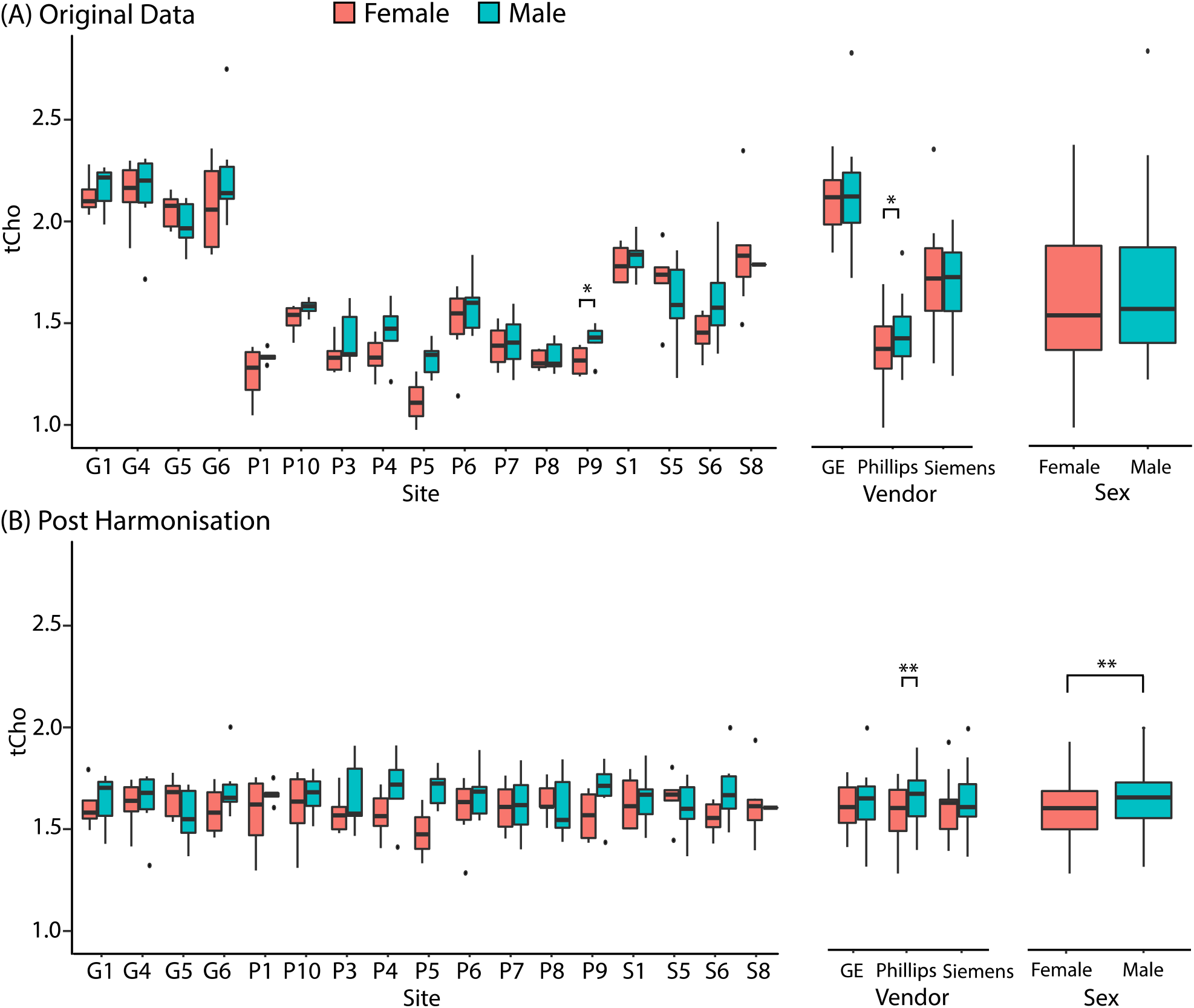
Sex differences in tCho levels (A) prior to and (B) following harmonisation. Box plots show median and interquartile range (IQR), dots represent outliers (more that 1.5xIQR).

To ensure that inclusion of the matrix of biological covariates did not introduce spurious biological variability, the harmonisation procedure was conducted with and without the matrix included. Both approaches yielded similar results and therefore results are not reported.

#### Creatine referenced data

Sex was significantly associated with tCho/tCr levels both prior to and following harmonisation. As with water referenced data, males had significantly higher tCho/tCr levels than females. No other biological variables were associated with creatine referenced metabolite levels prior to harmonisation. Following harmonisation, age was significantly associated with GABA+/tCr and Glx/tCr levels, with both metabolite levels decreasing with age (Figure 2, Table 3) see supplementary material for statistical model values.

#### Evaluation of sex effects

To confirm that ComBat harmonisation enabled detection of true biological effects, and to ensure the harmonisation procedure itself did not introduce false biological variability, the relationship between tCho (referenced to water) and sex was assessed within each site and vendor. This metabolite was chosen because tCho levels (referenced to water) were not significantly associated with sex prior to harmonisation, but were significantly associated with sex following harmonisation (Figure 3). As sex was significantly associated with tCho/tCr prior to and following harmonisation, this supports the interpretation that harmonisation revealed a true effect.

Prior to harmonisation, when analysing individual sites, only site P9 showed significant sex differences in tCho (males > females). Similar trends were found for sex-related differences within several other sites but did not reach statistical significance. When analysing vendor averaged data, a sex effect was only statistically significant for Phillips scanners. When averaging all data, there was no significant effect of sex. Following harmonisation, no individual sites showed significant effects of sex. Vendor averaged data showed a significant effect of sex in Phillips data only (males>females). When averaging all data, there was a significant effect of sex (males>females) (Figure 3).

## DISCUSSION

Here we show that ComBat harmonisation can effectively remove site and scanner variance in MRS data. We also show that removing this variance increased our ability to detect associations of sex and age with metabolite levels that were masked prior to harmonisation, confirming the utility of data harmonisation.

### Both vendor and site significantly effect metabolite levels

Linear modelling showed vendor and site to be significantly associated with metabolite levels when referencing to water, in agreement with previous studies of water referenced GABA+ and MM-sup GABA (Mikkelsen et al., 2019). No other studies have investigated these associations in water referenced PRESS data for the quantification of tNAA, tCr, tCho, Glx and Myo-Ins. Interestingly, vendor, but not site, was significantly associated with all metabolite levels when referencing to creatine. This is in contrast to Mikkelsen et al. (2017), who found that both vendor and site were significantly associated with GABA+/tCr levels and that site only was significantly associated with MM-sup GABA/tCr levels. Additionally, Považan et al., 2020 found significant effects of site on tNAA/tCr, tCho/tCr, Glx/tCr and Myo-Ins/tCr, with effects of vendor on tCho/tCr only. The previous studies included data from additional sites that were not available in the repository, which may have increased the site or vendor specific effects, and may explain the differences between the previous and current findings. Further, the current study used a generic pulse sequence to model the basis set (to emphasise effects of vendor in the data), whereas Považan et al., 2020 used vendor specific acquisition pulse waveforms to minimise vendor differences, which may have also caused differences in site or vendor specific effects.

### Harmonisation using ComBat successfully removes site and vendor effects

Following ComBat harmonisation by site, neither site nor vendor was associated with metabolite levels, whether referencing to water or to tCr, indicating ComBat harmonisation by site successfully removed this non-biological variance. Interestingly, harmonisation by vendor did not completely remove variability across different vendors. This could result from the nested nature of vendor within site, and variance from the two variables will be highly overlapped. We therefore recommend harmonisation by the lowest source of variance, in this case site.

### Harmonisation reveals previously masked biological effects

Harmonisation of MRS data revealed biological effects that were previously not detected, even when controlling for vendor and site in the model, indicating that (1) harmonisation with ComBat does remove non-biological variance and allows detection of biological effects and (2) controlling for these factors in statistical models is not enough to remove unwanted variance. These effects are consistent with the literature and confirm the utility of the harmonisation procedures.

Sex differences in tCho/tCr were seen both prior to and following harmonisation, but sex differences in water referenced tCho were only detected following data harmonisation across sites. Specifically, tCho/tCr and tCho referenced to water were significantly higher in males as compared to females. During the ComBat harmonisation procedure, inclusion of a matrix of biological variables is recommended to preserve biological variability. To ensure inclusion of this matrix did not introduce spurious biological variability, the harmonisation procedure was conducted with and without the matrix included, with no change in results. Therefore, we are confident the harmonisation procedure did not introduce spurious effects. Furthermore, this effect was not detected even when including site and vendor in the linear model as covariates, indicating that controlling for variables like site and vendor does not remove all the variance associated with these factors, and highlighting the usefulness of data harmonisation. tCho is generally considered to be a marker of cell density, and changes in tCho are thought to reflect alterations in membrane turnover (i.e. increase in membrane synthesis or breakdown) (Rae, 2014). However, evidence suggests that choline levels may also relate to acetylcholine function (Bell et al., 2018; Lindner et al., 2017). Our findings of higher tCho levels in males compared to females are in line with evidence from previous studies (Hadel et al., 2013; Považan et al., 2020) and may be due to the effects of estrogen. Craig et al., 2007 showed that tCho levels in women increased following 8 weeks of ovarian suppression, which significantly reduced estradiol levels. Thus, the higher level of tCho in males, who should have lower estrogen levels, is unsurprising.

Following harmonisation, we also found Glx/tCr and GABA+/tCr declined with increasing age, consistent with previous research (see Cichocka and Bereś, 2018 for a systematic review on metabolite changes and Porges et al., 2021 for a meta-analysis on GABA+ changes throughout the lifespan). Notably, prior adult studies examined age affects across a broader age range. This larger range facilitates the detection of smaller effects that are subtle in more restricted age ranges as in the current study (18-35). Indeed, the regression coefficients in the current study (−0.15 for Glx and −0.17 for GABA+) indicate a small effect size, which requires the increased power that is accomplished by the large number of participants included in this study. Additionally, neither of these effects were detectable prior to harmonisation, even when including vendor and site as covariates in the model, again highlighting the impact of non-biological variables and the benefits of data harmonisation.

### Conclusions

In conclusion, we show that ComBat is an effective approach to harmonise MRS data (both PRESS and GABA-edited MEGA-PRESS) and successfully removes non-biological variance due to site and vendor differences. Further, we confirm the utility of harmonisation by demonstrating that non-biological variance in MRS data can mask biological effects of interest and, upon harmonisation of data, expected biological effects of interest can be detected. We therefore recommend the use of ComBat harmonisation in multi-site MRS studies.

## Acknowledgements

This work was undertaken thanks in part to funding from: The Natural Sciences and Engineering Research Council of Canada, T. Chen Fong Postdoctoral Fellowship in Medical Imaging (TB), Harley N. Hotchkiss-Samuel Weiss Postdoctoral Fellowship (AW), Killam Postdoctoral Fellowship (AW), Harley N. Hotchkiss doctoral Fellowship (KG) and supported by the Hotchkiss Brain Institute and the Alberta Children’s Hospital Research Institute, University of Calgary. Data collection was undertaken thanks to NIH grant EB016089. ADH holds a Canada Research Chair in Magnetic Resonance Spectroscopy in Brain Injury and KOY holds a Ronald and Irene Ward Chair in Pediatric Brain Injury. The funders had no involvement in study design, in the collection, analysis, and interpretation of data, in writing the report, or in the decision to submit the paper for publication.

