## Supplemental Material for "Harmonisation of multi-site MRS data with ComBat" (from another version of this preprint: version 1, 2021-08-05)

Table 1: Summary of linear mixed-effects models for multi-level analysis of water referenced GABA+ data prior to harmonisation.

| Parameters | **(0a) Vendor only** | **(0b) Site only** | (1) Vendor and site | (4) Vendor, site and GMF | (5) Vendor, site and age | (6) Vendor, site and sex |
| --- | --- | --- | --- | --- | --- | --- |
| ***Regression Coefficients (fixed effects)*** |  |  |  |  |  |  |
| Intercept | **0.1144** | **0.0061** | 0.1125 | 0.1211 | 0.1132 | 0.1050 |
| GMF |  |  |  | 0.1179 |  |  |
| Age |  |  |  |  | -0.0416 |  |
| Sex |  |  |  |  |  | 0.0156 |
| ***Variance Components (random effects)*** |  |  |  |  |  |  |
| Vendor intercept | **0.7295** |  | 0.7175 | 0.3259 | 0.3718 | 0.2684 |
| Site intercept |  | **0.5452** | 0.1330 | 0.1573 | 0.1166 | 0.1260 |
| Vendor slope |  |  |  | 0.1501 | 0.0987 | 0.1335 |
| Site slope |  |  |  | 0.0711 | 0.0857 | 0.0858 |
| Vendor intercept x slope covariance |  |  |  | -0.3400 | -0.2400 | -0.3700 |
| Site intercept x slope covariance |  |  |  | -0.8900 | -1.000 | -0.9900 |
| Residual | **0.6050** | **0.4863** | 0.4854 | 0.4801 | 0.4852 | 0.4877 |
| ***Model fit outcome*** |  |  |  |  |  |  |
| Goodness-of-fit statistics (log lik) | **-260.5031** | **-256.7572** | -248.4910 | -248.4579 | -250.1976 | -249.8912 |
| ***ANOVA*** |  |  |  |  |  |  |
| Comparison | **0a vs 1** | **0b vs 1** |  | 1 vs 4 | 1 vs 5 | 0a vs 6 |
| p value | **9.56 x 10^-7^** | **1.32 x 10^-4^** |  | 0.5895 | 0.9891 | 1.0000 |

*GMF: Grey Matter Fraction*

Table 2: Summary of linear mixed-effects models for multi-level analysis of water referenced GABA+ data post harmonisation.

| Parameters | (0a) Vendor only | (0b) Site only | (1) Vendor and Site | **(4) GMF^†^** | (5) GMF and age^†^ | (6) GMF and sex^†^ |
| --- | --- | --- | --- | --- | --- | --- |
| ***Regression Coefficients (fixed effects)*** |  |  |  |  |  |  |
| Intercept | -1.83 x 10^-16^ | -1.83 x 10^-16^ | -1.83 x 10^-16^ | **-4.13 x 10^-16^** | -1.44 x 10^-16^ | 0.0075 |
| GMF |  |  |  | **0.1394** | 0.1589 | 0.1838 |
| Age |  |  |  |  | -0.0725 |  |
| Sex |  |  |  |  |  | -0.0154 |
| ***Variance Components (random effects)*** |  |  |  |  |  |  |
| Vendor intercept | 0 |  | 0 |  |  |  |
| Site intercept |  | 0 | 0 |  |  |  |
| Vendor slope |  |  |  |  |  |  |
| Site slope |  |  |  |  |  |  |
| Vendor intercept x slope covariance |  |  |  |  |  |  |
| Site intercept x slope covariance |  |  |  |  |  |  |
| Residual | 1 | 1 | 1 |  |  |  |
| ***Model fit outcome*** |  |  |  |  |  |  |
| Goodness-of-fit statistics (log lik) | -310.6019 | -310.6019 | -310.6019 | **-306.6897** | -304.567 | -305.0857 |
| ***ANOVA*** |  |  |  |  |  |  |
| Comparison | 0a vs 1 | 0b vs 1 |  |  |  |  |
| p value | 1.0000 | 1.0000 |  | **0.0398** | 0.3092 | 0.9081 |

*^†^ Fixed effect only model; GMF: Grey Matter Fraction*

Table 3: Summary of linear mixed-effects models for multi-level analysis of water referenced MS-GABA data prior to harmonisation.

| Parameters | **(0a) Vendor only** | **(0b) Site only** | (1) Vendor and site | (4) Vendor and GMF | (5) Vendor and age | (6) Vendor and sex |
| --- | --- | --- | --- | --- | --- | --- |
| ***Regression Coefficients (fixed effects)*** |  |  |  |  |  |  |
| Intercept | **-0.0417** | **-0.0463** | -0.0463 | -0.0422 | -0.0418 | -0.0122 |
| GMF |  |  |  | 0.0360 |  |  |
| Age |  |  |  |  | 0.0017 |  |
| Sex |  |  |  |  |  | -0.0632 |
| ***Variance Components (random effects)*** |  |  |  |  |  |  |
| Vendor intercept | **0.1778** |  | 7.34 x 10^-11^ | 0.0518 | 0.8345 | 1.0701 |
| Site Intercept |  | **0.6715** | 0.6715 |  |  |  |
| Vendor slope |  |  |  | 0.1695 | 1.3211 | 1.0842 |
| Site slope |  |  |  |  |  |  |
| Vendor intercept x slope covariance |  |  |  | -0.3900 | -0.96 | -0.92 |
| Site intercept x slope covariance |  |  |  |  |  |  |
| Residual | **0.898** | **0.3892** | 0.3892 | 0.9021 | 0.9023 | 0.9015 |
| ***Model fit outcome*** |  |  |  |  |  |  |
| Goodness-of-fit statistics (log lik) | **-289.2699** | **-226.461** | -226.4610 | -290.8137 | -291.0599 | -290.2593 |
| ***ANOVA*** |  |  |  |  |  |  |
| Comparison | **0a vs 1** | **0b vs 1** |  | 0a vs 4 | 0a vs 5 | 0a vs 6 |
| p value | **2.20 x 10^-16^** | **1.000** |  | 0.9563 | 1.000 | 0.9707 |

*GMF: Grey Matter Fraction*

Table 4: Summary of linear mixed-effects models for multi-level analysis of water referenced MS-GABA data post harmonisation.

| Parameters | (0a) Vendor only | (0b) Site only | (1) Vendor and Site | (4) GMF^†^ | (5) Age^†^ | (6) Sex^†^ |
| --- | --- | --- | --- | --- | --- | --- |
| ***Regression Coefficients (fixed effects)*** |  |  |  |  |  |  |
| Intercept | -3.59 x 10^-16^ | -3.59 x 10^-16^ | -3.59 x 10^-16^ | -5.23 x 10^-16^ | -3.51 x 10^-16^ | 0.0677 |
| GMF |  |  |  | 0.0565 |  |  |
| Age |  |  |  |  | -0.0136 |  |
| Sex |  |  |  |  |  | -0.1430 |
| ***Variance Components (random effects)*** |  |  |  |  |  |  |
| Vendor intercept | 0 |  | 0 |  |  |  |
| Site intercept |  | 0 | 0 |  |  |  |
| Vendor slope |  |  |  |  |  |  |
| Site slope |  |  |  |  |  |  |
| Vendor intercept x slope covariance |  |  |  |  |  |  |
| Site intercept x slope covariance |  |  |  |  |  |  |
| Residual | 1 | 1 | 1 |  |  |  |
| ***Model fit outcome*** |  |  |  |  |  |  |
| Goodness-of-fit statistics (log lik) | -297.8104 | -297.8104 | -297.8104 | -295.7231 | -296.0376 | -295.5204 |
| ***ANOVA*** |  |  |  |  |  |  |
| Comparison | 0a vs 1 | 0b vs 1 |  |  |  |  |
| p value | 1.000 | 1.0000 |  | 0.4170 | 0.8450 | 0.3030 |

*^†^ Fixed effect only model; GMF: Grey Matter Fraction*

Table 5: Summary of linear mixed-effects models for multi-level analysis of water referenced tNAA data prior to harmonisation.

| Parameters | **(0a) Vendor only** | **(0b) Site only** | (1) Vendor and site | (4) Vendor, site and GMF | (5) Vendor, site and age | (6) Vendor, site and sex |
| --- | --- | --- | --- | --- | --- | --- |
| ***Regression Coefficients (fixed effects)*** |  |  |  |  |  |  |
| Intercept | **0.3327** | **0.0265** | 0.3424 | 0.3410 | 0.3427 | 0.3478 |
| GMF |  |  |  | -0.0168 |  |  |
| Age |  |  |  |  | 0.0082 |  |
| Sex |  |  |  |  |  | -0.1126 |
| ***Variance Components (random effects)*** |  |  |  |  |  |  |
| Vendor intercept | **1.4240** |  | 1.4600 | 0.0605 | 0.0847 | 0.1324 |
| Site intercept |  | **1.0357** | 0.0436 | 0.0077 | 0.0798 | 0.0668 |
| Vendor slope |  |  |  | 0.0056 | 0.0096 | 0.0123 |
| Site slope |  |  |  | 0.0003 | 1.81 x 10^-4^ | 0.0013 |
| Vendor intercept x slope covariance | |  |  | 1.0000 | -0.2800 | -0.6800 |
| Site intercept x slope covariance | |  |  | -0.3800 | -0.6700 | -1.00 |
| Residual | **0.0976** | **0.0619** | 0.0619 | 0.0618 | 0.0621 | 0.0623 |
| ***Model fit outcome*** |  |  |  |  |  |  |
| Goodness-of-fit statistics (log lik) | **-57.1006** | **-49.4973** | -30.1377 | -32.7453 | -33.0573 | -32.4654 |
| ***ANOVA*** |  |  |  |  |  |  |
| Comparison | **0a vs 1** | **0b vs 1** |  | 1 vs 4 | 1 vs 5 | 1 vs 6 |
| p value | **2.12 x 10^-13^** | **1.29 x 10^-09^** |  | 0.9897 | 0.9994 | 0.9999 |

*GMF: Grey Matter Fraction*

Table 6: Summary of linear mixed-effects models for multi-level analysis of water referenced tNAA data post harmonisation.

| Parameters | (0a) Vendor only | (0b) Site only | (1) Vendor and Site | (4) GMF^†^ | (5) Age^†^ | (6) Sex^†^ |
| --- | --- | --- | --- | --- | --- | --- |
| ***Regression Coefficients (fixed effects)*** |  |  |  |  |  |  |
| Intercept | -1.34 x 10^-15^ | -1.34 x 10^-15^ | -1.34 x 10^-15^ | -1.52 x 10^-15^ | -1.34 x 10^-15^ | -0.0407 |
| GMF |  |  |  | -0.0725 |  |  |
| Age |  |  |  |  | 0.0389 |  |
| Sex |  |  |  |  |  | 0.0822 |
| ***Variance Components (random effects)*** |  |  |  |  |  |  |
| Vendor intercept | 0 |  | 0 |  |  |  |
| Site intercept |  | 0 | 0 |  |  |  |
| Vendor slope |  |  |  |  |  |  |
| Site slope |  |  |  |  |  |  |
| Vendor intercept x slope covariance | |  |  |  |  |  |
| Site intercept x slope covariance | |  |  |  |  |  |
| Residual | 1 | 1 | 1 |  |  |  |
| ***Model fit outcome*** |  |  |  |  |  |  |
| Goodness-of-fit statistics (log lik) | -267.9597 | -267.9597 | -267.9597 | -265.7633 | -266.1165 | -266.0933 |
| ***ANOVA*** |  |  |  |  |  |  |
| Comparison | 0a vs 1 | 0b vs 1 |  |  |  |  |
| p value | 1.0000 | 1.0000 |  | 0.3230 | 0.5960 | 0.5745 |

*^†^ Fixed effect only model;* *GMF: Grey Matter Fraction*

Table 7: Summary of linear mixed-effects models for multi-level analysis of water referenced tCr data prior to harmonisation.

| Parameters | **(0a) Vendor only** | **(0b) Site only** | (1) Vendor and site | (4) Vendor, site and GMF | (5) Vendor, site and age | (6) Vendor, site and sex |
| --- | --- | --- | --- | --- | --- | --- |
| ***Regression Coefficients (fixed effects)*** | **0.2909** | **0.3165** | 0.1603 | 0.3144 | 0.3114 | 0.3717 |
| Intercept |  |  |  | -0.0575 |  |  |
| GMF |  |  |  |  | 0.0488 |  |
| Age |  |  |  |  |  | -0.1284 |
| Sex |  |  |  |  |  |  |
| ***Variance Components (random effects)*** |  |  |  |  |  |  |
| Vendor Intercept | **1.0731** |  | 1.0894 | 0.0505 | 0.0314 | 0.0481 |
| Site Intercept |  | **0.9345** | 0.1857 | 0.2387 | 0.0873 | 1.27 x 10^-6^ |
| Vendor slope |  |  |  | 0.0122 | 0.0124 | 0.0158 |
| Site slope |  |  |  | 0.0004 | 0.0011 | 0.0019 |
| Vendor intercept x slope covariance | |  |  | -0.2400 | -0.3100 | -0.8600 |
| Site intercept x slope covariance | |  |  | -0.5200 | -0.0400 | 0.6100 |
| Residual | **0.3167** | **0.1604** | 0.1603 | 0.1594 | 0.15889 | 0.1573 |
| ***Model fit outcome*** |  |  |  |  |  |  |
| Goodness-of-fit statistics (log lik) | **-165.67733** | **-130.17013** | -121.06694 | -123.38596 | -122.42286 | -120.64356 |
| ***ANOVA*** |  |  |  |  |  |  |
| Comparison | **0a vs 1** | **0b vs 1** |  | 1 vs 4 | 1 vs 5 | 1 vs 6 |
| p value | **2.20 x 10^-16^** | **4.91 x 10^-5^** |  | 0.9997 | 0.7999 | 0.4580 |

*GMF: Grey Matter Fraction*

Table 8: Summary of linear mixed-effects models for multi-level analysis of water referenced tCr data post harmonisation.

| Parameters | (0a) Vendor only | (0b) Site only | | (1) Vendor and Site | | (4) GMF^†^ | | (5) Age^†^ | | (6) Sex^†^ |
| --- | --- | --- | --- | --- | --- | --- | --- | --- | --- | --- |
| ***Regression Coefficients (fixed effects)*** |  |  | |  | |  | |  | |  |
| Intercept | 6.06 x 10^-16^ | 6.06 x 10^-16^ | | 6.06 x 10^-16^ | | 4.32 x 10^-16^ | | 6.22 x 10^-16^ | | 0.0774 |
| GMF |  |  | |  | | 0.0931 | |  | |  |
| Age |  |  | |  | |  | | 0.1160 | |  |
| Sex |  |  | |  | |  | |  | | -0.1565 |
| ***Variance Components (random effects)*** |  |  | |  | |  | |  | |  |
| Vendor intercept | 0 |  | | 0 | |  | |  | |  |
| Site intercept |  | 0 | | 0 | |  | |  | |  |
| Vendor slope |  |  | |  | |  | |  | |  |
| Site slope |  |  | |  | |  | |  | |  |
| Vendor intercept x slope covariance | | |  | |  | |  | |  | |
| Site intercept x slope covariance | | |  | |  | |  | |  | |
| Residual | 1 | 1 | | 1 | |  | |  | |  |
| ***Model fit outcome*** |  |  | |  | |  | |  | |  |
| Goodness-of-fit statistics (log lik) | -267.9597 | -267.9597 | | -267.9597 | | -265.4408 | | -264.9863 | | -265.6787 |
| ***ANOVA*** |  |  | |  | |  | |  | |  |
| Comparison | 0a vs 1 | 0b vs 1 | |  | |  | |  | |  |
| p value | 1.0000 | 1.0000 | |  | | 0.204 | | 0.1130 | | 0.2850 |

*^†^ Fixed effect only model; GMF: Grey Matter Fraction*

Table 9: Summary of linear mixed-effects models for multi-level analysis of water referenced tCho data prior to harmonisation.

| Parameters | **(0a) Vendor only** | **(0b) Site only** | (1) Vendor and site | (4) Vendor, site and GMF | (5) Vendor, site and age | (6) Vendor, site and sex |
| --- | --- | --- | --- | --- | --- | --- |
| ***Regression Coefficients (fixed effects)*** |  |  |  |  |  |  |
| Intercept | **0.3083** | **0.0107** | 0.3093 | 0.3105 | 0.3101 | 0.2324 |
| GMF |  |  |  | 0.0143 |  |  |
| Age |  |  |  |  | 0.0211 |  |
| Sex |  |  |  |  |  | 0.1588 |
| ***Variance Components (random effects)*** |  |  |  |  |  |  |
| Vendor intercept | **1.1254** |  | 1.1380 | 0.2326 | 0.3090 | 0.1890 |
| Site intercept |  | **0.8779** | 0.0767 | 0.3218 | 0.0272 | 0.2879 |
| Vendor slope |  |  |  | 0.3157 | 0.24460 | 0.2807 |
| Site slope |  |  |  | 0.0328 | 0.0694 | 0.0231 |
| Vendor intercept x slope covariance | |  |  | 0.0800 | -0.9600 | -1.0000 |
| Site intercept x slope covariance | |  |  | -0.9900 | 0.2000 | 0.2700 |
| Residual | **0.2688** | **0.2054** | 0.2054 | 0.2063 | 0.2064 | 0.1990 |
| ***Model fit outcome*** |  |  |  |  |  |  |
| Goodness-of-fit statistics (log lik) | **-150.5826** | **-150.8943** | -137.1059 | -139.3423 | -139.3441 | -136.0997 |
| ***ANOVA*** |  |  |  |  |  |  |
| Comparison | **0a vs 1** | **0b vs 1** |  | 1 vs 4 | 1 vs 5 | 1 vs 6 |
| p value | **2.11 x 10^-7^** | **3.84 x 10^-7^** |  | 0.9997 | 0.9967 | 0.3472 |

*GMF: Grey Matter Fraction*

Table 10: Summary of linear mixed-effects models for multi-level analysis of water referenced tCho data post harmonisation.

| Parameters | (0a) Vendor only | (0b) Site only | (1) Vendor and Site | (4) GMF^†^ | (5) Age^†^ | **(6) Sex^†^** |
| --- | --- | --- | --- | --- | --- | --- |
| ***Regression Coefficients (fixed effects)*** |  |  |  |  |  |  |
| Intercept | -1.81 x 10^-16^ | -1.81 x 10^-16^ | -1.81 x 10^-16^ | -2.05 x 10^-16^ | -7.00 x 10^-17^ | **-0.1978** |
| GMF |  |  |  | 0.0317 |  |  |
| Age |  |  |  |  | 0.0848 |  |
| Sex |  |  |  |  |  | **0.3998** |
| ***Variance Components (random effects)*** |  |  |  |  |  |  |
| Vendor intercept | 0 |  | 0 |  |  |  |
| Site intercept |  | 0 | 0 |  |  |  |
| Vendor slope |  |  |  |  |  |  |
| Site slope |  |  |  |  |  |  |
| Vendor intercept x slope covariance | |  |  |  |  |  |
| Site intercept x slope covariance | |  |  |  |  |  |
| Residual | 1 | 1 | 1 |  |  |  |
| ***Model fit outcome*** |  |  |  |  |  |  |
| Goodness-of-fit statistics (log lik) | -267.9597 | -267.9597 | -267.9597 | -266.1647 | -265.5803 | **-262.4062** |
| ***ANOVA*** |  |  |  |  |  |  |
| Comparison | 0a vs 1 | 0b vs 1 |  |  |  |  |
| p value | 1.00 | 1 |  | 0.6660 | 0.2470 | **0.0058** |

*^†^ Fixed effect only model; GMF: Grey Matter Fraction*

Table 11: Summary of linear mixed-effects models for multi-level analysis of water referenced Glx data prior to harmonisation.

| Parameters | **(0a) Vendor only** | **(0b) Site only** | (1) Vendor and site | (4) Vendor, site and GMF | (5) Vendor, site and age | (6) Vendor, site and sex |
| --- | --- | --- | --- | --- | --- | --- |
| ***Regression Coefficients (Fixed Effects)*** | |  |  |  |  |  |
| Intercept | **0.2982** | **0.0204** | 0.3028 | 0.3083 | 0.3020 | 0.3174 |
| GMF |  |  |  | 0.0564 |  |  |
| Age |  |  |  |  | -0.0211 |  |
| Sex |  |  |  |  |  | -0.0302 |
| ***Variance Components (Random Effects)*** | |  |  |  |  |  |
| Vendor intercept | **1.0486** |  | 1.0688 | 0.2549 | 0.2158 | 0.2739 |
| Site intercept |  | **0.8392** | 0.0917 | 0.1926 | 0.1819 | 0.2246 |
| Vendor slope |  |  |  | 0.0062 | 0.0063 | 0.0038 |
| Site slope |  |  |  | 0.0004 | 0.0020 | 0.0006 |
| Vendor intercept x slope covariance |  |  |  | -0.6300 | -0.6700 | -0.2000 |
| Site intercept x slope covariance |  |  |  | -0.7200 | -1.0000 | -0.7700 |
| Residual | **0.3178** | **0.2432** | 0.2433 | 0.2423 | 0.2440 | 0.2443 |
| ***Model fit outcome*** |  |  |  |  |  |  |
| Goodness-of-fit statistics (log lik) | **-165.9849** | **-165.0126** | -165.0126 | -154.1309 | -154.9410 | -154.3659 |
| ***ANOVA*** |  |  |  |  |  |  |
| Comparison | **0a vs 1** | **0b vs 1** |  | 1 vs 4 | 1 vs 5 | 1 vs 6 |
| p value | **2.28 x 10^-7^** | **1.87 x 10^-6^** |  | 1.8906 | 0.9977 | 0.9994 |

*GMF: Grey Matter Fraction*

Table 12: Summary of linear mixed-effects models for multi-level analysis of water referenced Glx data post harmonisation.

| Parameters | (0a) Vendor only | (0b) Site only | (1) Vendor and site | (4) GMF^†^ | (5) Age^†^ | (6) Sex^†^ |
| --- | --- | --- | --- | --- | --- | --- |
| ***Regression Coefficients (fixed effects)*** |  |  |  |  |  |  |
| Intercept | 3.42 x 10^-16^ | 3.42 x 10^-16^ | 3.42 x 10^-16^ | 5.40 x 10^-16^ | 3.81 x 10^-16^ | 0.0085 |
| GMF |  |  |  | 0.1186 |  |  |
| Age |  |  |  |  | -0.0626 |  |
| Sex |  |  |  |  |  | -0.0173 |
| ***Variance Components (random effects)*** |  |  |  |  |  |  |
| Vendor intercept | 0 |  | 0 |  |  |  |
| Site intercept |  | 0 | 0 |  |  |  |
| Vendor slope |  |  |  |  |  |  |
| Site slope |  |  |  |  |  |  |
| Vendor intercept x slope covariance |  |  |  |  |  |  |
| Site intercept x slope covariance |  |  |  |  |  |  |
| Residual | 1 | 1 | 1 |  |  |  |
| **Model fit outcome** |  |  |  |  |  |  |
| Goodness-of-fit statistics (log lik) | -267.9597 | -267.9597 | -267.9597 | -264.9266 | -265.8905 | -266.2521 |
| ***ANOVA*** |  |  |  |  |  |  |
| Comparison | 0a vs 1 | 0b vs 1 |  |  |  |  |
| p value | 1.00 | 1 |  | 0.1050 | 0.3940 | 0.9060 |

*^†^ Fixed effect only model; GMF: Grey Matter Fraction*

Table 13: Summary of linear mixed-effects models for multi-level analysis of water referenced Myo-Ins data prior to harmonisation.

| Parameters | **(0a) Vendor only** | **(0b) Site only** | (1) Vendor and site | (4) Vendor, site and GMF | (5) Vendor, site and age | (6) Vendor, site and sex |
| --- | --- | --- | --- | --- | --- | --- |
| ***Regression Coefficients (fixed effects)*** |  |  |  |  |  |  |
| Intercept | **0.1815** | **0.0083** | 0.1812 | 0.1836 | 0.18079 | 0.18208 |
| GMF |  |  |  | -0.0288 |  |  |
| Age |  |  |  |  | -0.01168 |  |
| Sex |  |  |  |  |  | -0.001749 |
| ***Variance Components (random effects)*** |  |  |  |  |  |  |
| Vendor Intercept | **1.3102** |  | 1.3816 | 0.0572 | 0.1978 | 0.3369 |
| Site Intercept |  | **0.9111** | 0.1334 | 0.3237 | 0.1649 | 0.2039 |
| Vendor slope |  |  |  | 0.0308 | 0.0190 | 0.0092 |
| Site slope |  |  |  | 0.0044 | 0.0018 | 0.0011 |
| Vendor intercept x slope covariance | |  |  | -0.2300 | 0.3500 | 0.8100 |
| Site intercept x slope covariance | |  |  | 0.7700 | -0.4900 | -0.5900 |
| Residual | **0.3242** | **0.2156** | 0.2157 | 0.2163 | 0.2169 | 0.2169 |
| ***Model fit outcome*** |  |  |  |  |  |  |
| Goodness-of-fit statistics (log lik) | **-168.0496** | **-155.3348** | -144.7690 | -146.7921 | -147.1006 | -146.5183 |
| ***ANOVA*** |  |  |  |  |  |  |
| Comparison | **0a vs 1** | **0b vs 1** |  | 1 vs 4 | 1 vs 5 | 1 vs 6 |
| p value | **9.20 x 10^-12^** | **1.20 x 10^-5^** |  | 0.9925 | 0.9998 | 1.0000 |

*GMF: Grey Matter Fraction*

Table 14: Summary of linear mixed-effects models for multi-level analysis of water referenced Myo-Ins data post harmonisation.

| Parameters | (0a) Vendor only | (0b) Site only | (1) Vendor and Site | (4) GMF^†^ | (5) Age^†^ | (6) Sex^†^ |
| --- | --- | --- | --- | --- | --- | --- |
| ***Regression Coefficients (fixed effects)*** |  |  |  |  |  |  |
| Intercept | -3.40 x 10^-16^ | -3.40 x 10^-16^ | -3.40 x 10^-16^ | -3.73 x 10^-16^ | -3.85 x 10^-16^ | 0.0290 |
| GMF |  |  |  | 0.0408 |  |  |
| Age |  |  |  |  | -0.0321 |  |
| Sex |  |  |  |  |  | -0.0587 |
| ***Variance Components (random effects)*** |  |  |  |  |  |  |
| Vendor Intercept | 0 |  | 0 |  |  |  |
| Site Intercept |  | 0 | 0 |  |  |  |
| Vendor slope |  |  |  |  |  |  |
| Site slope |  |  |  |  |  |  |
| Vendor intercept x slope covariance | |  |  |  |  |  |
| Site intercept x slope covariance | |  |  |  |  |  |
| Residual | 1 | 1 | 1 |  |  |  |
| ***Model fit outcome*** |  |  |  |  |  |  |
| Goodness-of-fit statistics (log lik) | -267.9597 | -267.9597 | -267.9597 | -266.1028 | -266.1625 | -266.177 |
| ***ANOVA*** |  |  |  |  |  |  |
| Comparison | 0a vs 1 | 0b vs 1 |  |  |  |  |
| p value | 1.0000 | 1.0000 |  | 0.5787 | 0.6620 | 0.6890 |

*^†^ Fixed effect only model; GMF: Grey Matter Fraction*

Table 15: Summary of linear mixed-effects models for multi-level analysis of GABA+/tCr data prior to harmonisation.

| Parameters | **(0a) Vendor only** | **(0b) Site only** | | (1) Vendor and site | | (4) Vendor and GMF | | (5) Vendor and age | | (6) Vendor and sex | |
| --- | --- | --- | --- | --- | --- | --- | --- | --- | --- | --- | --- |
| ***Regression Coefficients (fixed effects)*** |  |  | |  | |  | |  | |  | |
| Intercept (B0) | **0.0480** | **-0.0458** | | 0.0040 | | 0.0293 | | 0.0487 | | 0.0014 | |
| GMF |  |  | |  | | -0.0433 | |  | |  | |
| Age |  |  | |  | |  | | -0.0172 | |  | |
| Sex |  |  | |  | |  | |  | | 0.0978 | |
| ***Variance Components (random effects)*** |  |  | |  | |  | |  | |  | |
| Vendor intercept | **0.2641** |  | | 0.2989 | | 0.5577 | | 0.9337 | | 0.7758 | |
| Site intercept |  | **0.4301** | | 0.2021 | |  | |  | |  | |
| Vendor slope |  |  | |  | | 4.1405 | | 18.3149 | | 12.1195 | |
| Site slope |  |  | |  | |  | |  | |  | |
| Vendor intercept x slope covariance | | |  | |  | | -0.9100 | | -0.9600 | | -0.9600 |
| Site intercept x slope covariance | | |  | |  | |  | |  | |  |
| Residual | **0.8491** | **0.6233** | | 0.6240 | | 0.8239 | | 0.8532 | | 0.8512 | |
| ***Model fit outcome*** |  |  | |  | |  | |  | |  | |
| Goodness-of-fit statistics (log lik) | **-271.7719** | **-256.8059** | | -267.0776 | | -294.1040 | | -273.5326 | | -272.6062 | |
| ***ANOVA*** |  |  | |  | |  | |  | |  | |
| Comparison | **0a vs 1** | **0b vs 1** | |  | | 0a vs 4 | | 0a vs 5 | | 0a vs 6 | |
| p value | **1.23 x 10^-8^** | **0.2400** | |  | | 0.9491 | | 0.9900 | | 0.9000 | |

*GMF: Grey Matter Fraction*

Table 16: Summary of linear mixed-effects models for multi-level analysis of GABA+/tCr data post harmonisation.

| Parameters | (0a) Vendor only | (0b) Site only | (1) Vendor and Site | (4) GMF^†^ | (5) Age^†^ | (6) Age and Sex^†^ |
| --- | --- | --- | --- | --- | --- | --- |
| ***Regression Coefficients (fixed effects)*** |  |  |  |  |  |  |
| Intercept | 5.42 x 10^-16^ | 5.42 x 10^-16^ | 5.42 x 10^-16^ | 5.66 x 10^-16^ | 6.07 x 10^-16^ | -0.0401 |
| GMF |  |  |  | 0.0049 |  |  |
| Age |  |  |  |  | -0.1732 |  |
| Sex |  |  |  |  |  | 0.0825 |
| ***Variance Components (random effects)*** |  |  |  |  |  |  |
| Vendor intercept | 0 | 0 | 0 |  |  |  |
| Site intercept |  |  | 0 |  |  |  |
| Vendor slope |  |  |  |  |  |  |
| Site slope |  |  |  |  |  |  |
| Vendor intercept x slope covariance | |  |  |  |  |  |
| Site intercept x slope covariance | |  |  |  |  |  |
| Residual | 1 | 1 | 1 |  |  |  |
| ***Model fit outcome*** |  |  |  |  |  |  |
| Goodness-of-fit statistics (log lik) | -310.6019 | -310.6019 | -310.6019 | -308.8249 | -307.1174 | -306.9797 |
| ***ANOVA*** |  |  |  |  |  |  |
| Comparison | 0a vs 1 | 0b vs 1 |  |  |  |  |
| p value | 1.0000 | 1.0000 |  | 0.9430 | 0.0104 | 0.5440 |

*^†^ Fixed effect only model; GMF: Grey Matter Fraction*

Table 17: Summary of linear mixed-effects models for multi-level analysis of MS-GABA/tCr data prior to harmonisation.

| Parameters | **(0a) Vendor only** | | (0b) Site only | | (1) Vendor and site | | (4) Vendor and GMF | | (5) Vendor and age | | (6) Vendor and sex |
| --- | --- | --- | --- | --- | --- | --- | --- | --- | --- | --- | --- |
| ***Regression Coefficients (fixed effects)*** |  | |  | |  | |  | |  | |  |
| Intercept | **-0.0395** | | -0.0384 | | -0.0384 | | -0.0421 | | -0.0399 | | -0.0216 |
| GMF |  | |  | |  | | -0.0660 | |  | |  |
| Age |  | |  | |  | |  | | 0.0087 | |  |
| Sex |  | |  | |  | |  | |  | | -0.0387 |
| ***Variance Components (random effects)*** |  | |  | |  | |  | |  | |  |
| Vendor intercept | **0.0463** | |  | | 0.0000 | | 0.0523 | | 0.0446 | | 0.0444 |
| Site intercept |  | | 0.6334 | | 0.6334 | |  | |  | |  |
| Vendor slope |  | |  | |  | | 1.1079 | | 1.0992 | | 1.1075 |
| Site slope |  | |  | |  | |  | |  | |  |
| Vendor intercept x slope covariance |  | |  | |  | | 0.0200 | |  | |  |
| Site intercept x slope covariance |  | |  | |  | |  | |  | |  |
| Residual | **0.9668** | | 0.4536 | | 0.4536 | | 0.9669 | | 0.9713 | | 0.9710 |
| ***Model fit outcome*** |  | |  | |  | |  | |  | |  |
| Goodness-of-fit statistics (log lik) | **-295.7299** | | -240.6018 | | -240.6018 | | -297.0163 | | -297.4774 | | -296.7574 |
| ***ANOVA*** |  | |  | |  | |  | |  | |  |
| Comparison | **0a vs 1** | | 0b vs 1 | |  | | 0a vs 4 | | 0a vs 5 | | 0a vs 6 |
| p value | **p < 0.0001** | | 1.0000 | |  | | 1.0000 | | 0.9999 | | 0.9954 |

*GMF: Grey Matter Fraction*

Table 18: Summary of linear mixed-effects models for multi-level analysis of MS-GABA/tCr data post harmonisation.

| Parameters | (0a) Vendor only | (0b) Site only | (1) Vendor and Site | (4) GMF^†^ | (5) Age^†^ | (6) Sex^†^ |
| --- | --- | --- | --- | --- | --- | --- |
| ***Regression Coefficients (fixed effects)*** |  |  |  |  |  |  |
| Intercept | 2.61 x 10^-16^ | 2.61 x 10^-16^ | 2.61 x 10^-16^ | 8.37 x 10^-17^ | 2.44 x 10^-16^ | 0.0741 |
| GMF |  |  |  | 0.0169 |  |  |
| Age |  |  |  |  | -0.0116 |  |
| Sex |  |  |  |  |  | -0.1565 |
| ***Variance Components (random effects)*** |  |  |  |  |  |  |
| Vendor intercept | 0 |  | 0 |  |  |  |
| Site intercept |  | 0 | 0 |  |  |  |
| Vendor slope |  |  |  |  |  |  |
| Site slope |  |  |  |  |  |  |
| Vendor intercept x slope covariance |  |  |  |  |  |  |
| Site intercept x slope covariance |  |  |  |  |  |  |
| Residual | 1 | 1 | 1 |  |  |  |
| ***Model fit outcome*** |  |  |  |  |  |  |
| Goodness-of-fit statistics (log lik) | -297.8104 | -297.8104 | -297.8104 | -296.0270 | -296.0429 | -295.4141 |
| ***ANOVA*** |  |  |  |  |  |  |
| Comparison | 0a vs 1 | 0b vs 1 |  |  |  |  |
| p value | 1.0000 | 1.0000 |  | 0.8080 | 0.8680 | 0.2600 |

*^†^ Fixed effect only model; GMF: Grey Matter Fraction*

Table 19: Summary of linear mixed-effects models for multi-level analysis of tNAA/tCr data prior to harmonisation.

| Parameters | **(0a) Vendor only** | (0b) Site only | | (1) Vendor and site | | (4) Vendor and GMF | | (5) Vendor and age | | (6) Vendor and sex | |
| --- | --- | --- | --- | --- | --- | --- | --- | --- | --- | --- | --- |
| ***Regression Coefficients (fixed effects)*** |  |  | |  | |  | |  | |  | |
| Intercept | **0.0978** | -0.0053 | | 0.0290 | | 0.0840 | | 0.0962 | | -0.0480 | |
| GMF |  |  | |  | | -0.1298 | |  | |  | |
| Age |  |  | |  | |  | | -0.0890 | |  | |
| Sex |  |  | |  | |  | |  | | 0.3002 | |
| ***Variance Components (random effects)*** |  |  | |  | |  | |  | |  | |
| Vendor intercept | **0.1038** |  | | 0.0396 | | 0.1817 | | 0.1875 | | 0.1999 | |
| Site intercept |  | 0.4219 | | 0.3953 | |  | |  | |  | |
| Vendor slope |  |  | |  | | 0.2987 | | 0.3222 | | 0.3431 | |
| Site slope |  |  | |  | |  | |  | |  | |
| Vendor intercept x slope covariance | | |  | |  | | -0.9900 | | -1.0000 | | -1.0000 |
| Site intercept x slope covariance | | |  | |  | |  | |  | |  |
| Residual | **0.9288** | 0.5924 | | 0.5924 | | 0.9196 | | 0.9262 | | 0.9108 | |
| ***Model fit outcome*** |  |  | |  | |  | |  | |  | |
| Goodness-of-fit statistics (log lik) | **-263.0275** | -236.3948 | | -236.3106 | | -263.1736 | | -263.9660 | | -261.7811 | |
| ***ANOVA*** |  |  | |  | |  | |  | |  | |
| Comparison | **0a vs 1** | 0b vs 1 | |  | | 0a vs 4 | | 0a vs 5 | | 0a vs 6 | |
| p value | **2.65 x 10^-13^** | 1.0000 | |  | | 0.3583 | | 0.6534 | | 0.2051 | |

*GMF: Grey Matter Fraction*

Table 20: Summary of linear mixed-effects models for multi-level analysis of tNAA/tCr data post harmonisation.

| Parameters | (0a) Vendor only | (0b) Site only | (1) Vendor and Site | (4) GMF^†^ | (5) Age^†^ | (6) Sex^†^ |
| --- | --- | --- | --- | --- | --- | --- |
| ***Regression Coefficients (fixed effects)*** |  |  |  |  |  |  |
| Intercept | 4.23 x 10^-16^ | 4.23 x 10^-16^ | 4.23 x 10^-16^ | 6.94 x 10^-16^ | 4.22 x 10^-16^ | -0.0877 |
| GMF |  |  |  | -0.1376 |  |  |
| Age |  |  |  |  | -0.0835 |  |
| Sex |  |  |  |  |  | 0.1772 |
| ***Variance Components (random effects)*** |  |  |  |  |  |  |
| Vendor intercept | 0 |  | 0 |  |  |  |
| Site intercept |  | 0 | 0 |  |  |  |
| Vendor slope |  |  |  |  |  |  |
| Site slope |  |  |  |  |  |  |
| Vendor intercept x slope covariance | |  |  |  |  |  |
| Site intercept x slope covariance | |  |  |  |  |  |
| Residual | 1 | 1 | 1 |  |  |  |
| ***Model fit outcome*** |  |  |  |  |  |  |
| Goodness-of-fit statistics (log lik) | -267.9597 | -267.9597 | -267.9597 | -264.4634 | -265.6011 | -265.5139 |
| ***ANOVA*** |  |  |  |  |  |  |
| Comparison | 0a vs 1 | 0b vs 1 |  |  |  |  |
| p value | 1.0000 | 1.0000 |  | 0.0598 | 0.2540 | 0.2250 |

*^†^ Fixed effect only model; GMF: Grey Matter Fraction*

Table 21: Summary of linear mixed-effects models for multi-level analysis of tCho/tCr data prior to harmonisation.

| Parameters | **(0a) Vendor only** | (0b) Site only | | (1) Vendor and site | | (4) Vendor and GMF | | (5) Vendor and age | | (6) Vendor and sex | |
| --- | --- | --- | --- | --- | --- | --- | --- | --- | --- | --- | --- |
| ***Regression Coefficients (fixed effects)*** |  |  | |  | |  | |  | |  | |
| Intercept | **0.1139** | -0.0234 | | 0.0691 | | 0.0947 | | 0.1145 | | -0.1481 | |
| GMF |  |  | |  | | -0.1743 | |  | |  | |
| Age |  |  | |  | |  | | 0.0363 | |  | |
| Sex |  |  | |  | |  | |  | | 0.5411 | |
| ***Variance Components (random effects)*** |  |  | |  | |  | |  | |  | |
| Vendor intercept | **0.1245** |  | | 0.105 | | 0.0787 | | 0.1050 | | 0.1163 | |
| Site intercept |  | 0.2673 | | 0.1896 | |  | |  | |  | |
| Vendor slope |  |  | |  | | 1.1081 | | 1.2183 | | 1.1115 | |
| Site slope |  |  | |  | |  | |  | |  | |
| Vendor intercept x slope 1 covariance | | |  | |  | | -0.1000 | | -0.1100 | | -0.1300 |
| Site intercept x slope 1 covariance | | |  | |  | |  | |  | |  |
| Residual (participant) | **0.9112** | 0.7562 | | 0.7565 | | 0.8903 | | 0.9147 | | 0.8414 | |
| ***Model fit outcome*** |  |  | |  | |  | |  | |  | |
| Goodness-of-fit statistics (log lik) | **-261.4174** | -254.4987 | | -253.5488 | | -260.2769 | | -263.0231 | | -254.6699 | |
| ***ANOVA*** |  |  | |  | |  | |  | |  | |
| Comparison | **0a vs 1** | 0b vs 1 | |  | | 0a vs 4 | | 0a vs 5 | | 0a vs 6 | |
| p value | **7.70 x 10^-5^** | 0.2934 | |  | | 0.1162 | | 0.9676 | | 0.0013 | |

*GMF: Grey Matter Fraction*

Table 22: Summary of linear mixed-effects models for multi-level analysis of tCho/tCr data post harmonisation.

| Parameters | (0a) Vendor only | (0b) Site only | (1) Vendor and Site | (4) GMF^†^ | (5) Age^†^ | **(6) Sex^†^** |
| --- | --- | --- | --- | --- | --- | --- |
| ***Regression Coefficients (fixed effects)*** |  |  |  |  |  |  |
| Intercept | -9.36 x 10^-17^ | -9.36 x 10^-17^ | -9.36 x 10^-17^ | -3.11 x 10^-16^ | -1.16 x 10^-16^ | **-0.3047** |
| GMF |  |  |  | -0.0412 |  |  |
| Age |  |  |  |  | -0.0523 |  |
| Sex |  |  |  |  |  | **0.6160** |
| ***Variance Components (random effects)*** |  |  |  |  |  |  |
| Vendor intercept | 0 |  | 0 |  |  |  |
| Site intercept |  | 0 | 0 |  |  |  |
| Vendor slope |  |  |  |  |  |  |
| Site slope |  |  |  |  |  |  |
| Vendor intercept x slope covariance | |  |  |  |  |  |
| Site intercept x slope covariance | |  |  |  |  |  |
| Residual | 1 | 1 | 1 |  |  |  |
| ***Model fit outcome*** |  |  |  |  |  |  |
| Goodness-of-fit statistics (log lik) | -267.9597 | -267.9597 | -267.9597 | -266.0994 | -266.0017 | **-256.8372** |
| ***ANOVA*** |  |  |  |  |  |  |
| Comparison | 0a vs 1 | 0b vs 1 |  |  |  |  |
| p value | 1.0000 | 1.0000 |  | 0.5750 | 0.4760 | **1.62 x 10^-5^** |

*^†^ Fixed effect only model; GMF: Grey Matter Fraction*

Table 23: Summary of linear mixed-effects models for multi-level analysis of Glx/tCr data prior to harmonisation.

| Parameters | **(0a) Vendor only** | (0b) Site only | | (1) Vendor and site | (4) Vendor and GMF | (5) Vendor and age | (6) Vendor and sex |
| --- | --- | --- | --- | --- | --- | --- | --- |
| ***Regression Coefficients (fixed effects)*** |  |  | |  |  |  |  |
| Intercept | **0.1410** | -0.0033 | | -0.0962 | 0.1354 | 0.1405 | 0.0106 |
| GMF |  |  | |  | -0.0503 |  |  |
| Age |  |  | |  |  | -0.0257 |  |
| Sex |  |  | |  |  |  | 0.2685 |
| ***Variance Components (random effects)*** |  |  | |  |  |  |  |
| Vendor intercept | **0.1900** |  | | 0.1455 | 0.1440 | 0.1710 | 0.1232 |
| Site intercept |  | 0.4346 | | 0.3306 |  |  |  |
| Vendor slope |  |  | |  | 0.2367 | 0.2590 | 0.2233 |
| Site slope |  |  | |  |  |  |  |
| Vendor intercept x slope covariance | |  |  | | -1.0000 | -0.9900 | -0.9900 |
| Site intercept x slope covariance | |  |  | |  |  |  |
| Residual | **0.8656** | 0.5932 | | 0.5933 | 0.8684 | 0.8697 | 0.8518 |
| ***Model fit outcome*** |  |  | |  |  |  |  |
| Goodness-of-fit statistics (log lik) | **-257.037** | -236.7208 | | --235.8876 | -258.5079 | -258.7317 | -256.1539 |
| ***ANOVA*** |  |  | |  |  |  |  |
| Comparison | **0a vs 1** | 0b vs 1 | |  | 0a vs 4 | 0a vs 5 | 0a vs 6 |
| p value | **8.15 x 10^-11^** | 0.3428 | |  | 0.3583 | 0.9853 | 0.2698 |

*GMF: Grey Matter Fraction*

Table 24: Summary of linear mixed-effects models for multi-level analysis of Glx/tCr data post harmonisation.

| Parameters | (0a) Vendor only | (0b) Site only | (1) Vendor and Site | (4) GMF^†^ | **(5) Age^†^** | (6) Age and Sex^†^ |
| --- | --- | --- | --- | --- | --- | --- |
| ***Regression Coefficients (fixed effects)*** |  |  |  |  |  |  |
| Intercept | 4.19 x 10^-16^ | 4.19 x 10^-16^ | 4.19 x 10^-16^ | -5.48 x 10^-16^ | **3.86 x 10^-16^** | -0.0941 |
| GMF |  |  |  | -0.0153 |  | -0.1647 |
| Age |  |  |  |  | **-0.1470** | 0.1902 |
| Sex |  |  |  |  |  |  |
| ***Variance Components (random effects)*** |  |  |  |  |  |  |
| Vendor intercept | 0 |  | 0 |  |  |  |
| Site intercept |  | 0 | 0 |  |  |  |
| Vendor slope |  |  |  |  |  |  |
| Site slope |  |  |  |  |  |  |
| Vendor intercept x slope covariance | |  |  |  |  |  |
| Site intercept x slope covariance | |  |  |  |  |  |
| Residual | 1 | 1 | 1 |  |  |  |
| ***Model fit outcome*** |  |  |  |  |  |  |
| Goodness-of-fit statistics (log lik) | -267.9597 | -267.9597 | -267.9597 | -266.2370 | **-264.2047** | -263.3575 |
| ***ANOVA*** |  |  |  |  |  |  |
| Comparison | 0a vs 1 | 0b vs 1 |  |  |  |  |
| p value | 1.0000 | 1.0000 |  | 0.8350 | **0.0441** | 0.1972 |

*^†^ Fixed effect only model; GMF: Grey Matter Fraction*

Table 25: Summary of linear mixed-effects models for multi-level analysis of Myo-Ins/tCr data prior to harmonisation.

| Parameters | **(0a) Vendor only** | (0b) Site only | (1) Vendor and site | (4) Vendor and GMF | (5) Vendor and age | (6) Vendor and sex |
| --- | --- | --- | --- | --- | --- | --- |
| ***Regression Coefficients (fixed effects)*** |  |  |  |  |  |  |
| Intercept | **-0.0955** | -0.0279 | -0.1126 | -0.1057 | -0.0965 | -0.1998 |
| GMF |  |  |  | -0.0942 |  |  |
| Age |  |  |  |  | -0.0716 |  |
| Sex |  |  |  |  |  | 0.2146 |
| ***Variance Components (random effects)*** |  |  |  |  |  |  |
| Vendor intercept | **0.4882** |  | 0.465 | 1.2228 | 0.8109 | 0.8302 |
| Site intercept |  | 0.684 | 0.4236 |  |  |  |
| Vendor slope |  |  |  | 2.6453 | 1.3728 | 1.3861 |
| Site slope |  |  |  |  |  |  |
| Vendor intercept x slope covariance | |  |  | -0.7900 | -0.6800 | -0.6900 |
| Site intercept x slope covariance | |  |  |  |  |  |
| Residual | **0.7474** | 0.4021 | 0.4021 | 0.7426 | 0.7463 | 0.7399 |
| ***Model fit outcome*** |  |  |  |  |  |  |
| Goodness-of-fit statistics (log lik) | **-244.3448** | -206.5827 | -204.7978 | -245.1410 | -245.5465 | -244.0486 |
| ***ANOVA*** |  |  |  |  |  |  |
| Comparison | **0a vs 1** | 0b vs 1 |  | 0a vs 4 | 0a vs 5 | 0a vs 6 |
| p value | **2.20 x 10^-16^** | 0.1297 |  | 0.5946 | 0.7308 | 0.4022 |

*GMF: Grey Matter Fraction*

Table 26: Summary of linear mixed-effects models for multi-level analysis of Myo-Ins/tCr data post harmonisation.

| Parameters | (0a) Vendor only | (0b) Site only | (1) Vendor and Site | (4) GMF^†^ | (5) Age^†^ | (6) Sex^†^ |
| --- | --- | --- | --- | --- | --- | --- |
| ***Regression Coefficients (fixed effects)*** |  |  |  |  |  |  |
| Intercept | 5.74 x 10^-16^ | 5.74 x 10^-16^ | 5.74 x 10^-16^ | 4.97 x 10^-17^ | 5.57 x 10^-16^ | -0.1171 |
| GMF |  |  |  | -0.0411 |  |  |
| Age |  |  |  |  | -0.1322 |  |
| Sex |  |  |  |  |  | 0.2367 |
| ***Variance Components (random effects)*** |  |  |  |  |  |  |
| Vendor intercept | 0 |  | 0 |  |  |  |
| Site intercept |  | 0 | 0 |  |  |  |
| Vendor slope |  |  |  |  |  |  |
| Site slope |  |  |  |  |  |  |
| Vendor intercept x slope covariance | |  |  |  |  |  |
| Site intercept x slope covariance | |  |  |  |  |  |
| Residual | 1 | 1 | 1 |  |  |  |
| ***Model fit outcome*** |  |  |  |  |  |  |
| Goodness-of-fit statistics (log lik) | -267.9597 | -267.9597 | -267.9597 | -265.1003 | -264.6029 | -264.9264 |
| ***ANOVA*** |  |  |  |  |  |  |
| Comparison | 0a vs 1 | 0b vs 1 |  |  |  |  |
| p value | 1.0000 | 1.0000 |  | 0.5760 | 0.0706 | 0.1050 |

*^†^ Fixed effect only model; GMF: Grey Matter Fraction*
